# Forty-Eight Hour Conditioning Produces a Robust Long Lasting Flavor Preference in Rats

**DOI:** 10.1101/259804

**Authors:** Adam Kimbrough, Thomas A. Houpt

**Affiliations:** Department of Biological Science, Program in Neuroscience, Florida State University, Tallahassee, Fl; The Scripps Research Institute, Neuroscience Department, La Jolla, Ca

**Author notes:** Correspondence: Thomas A. Houpt, Department of Biological Science, 3010 King Building, The Florida State University, Tallahassee, FL 32306–4295, USA.

**Keywords:** Flavor, learning, glucose, flavor preference

## Abstract

Conditioned flavor preference (CFP) learning is a form of associative learning in ingestive behavior. CFP Learning can be rapid and produces preferences of varying strengths that can be exceptionally persistent. We sought to establish a method to produce a robust long-lasting CFP in rats. Rats were given 48-h access (conditioning) to a CS+ flavor (grape or cherry 0.05% Kool-Aid, counterbalanced) mixed with 8% glucose and 0.05% saccharin. In order to determine the strength of conditioning rats were given 14 consecutive days of 24-h access to CS+ and CS- flavors mixed only with 0.05% Kool-Aid and 0.05% saccharin (extinction), then further tested 34 days after the last extinction test (48 days post conditioning) for 2 consecutive days with the CS+ and CS-. We found that not only did the learned CFP fail to extinguish over 14 days of testing, but it also persisted for at least 48 days after conditioning. These data provide a method to produce a robust, long lasting and persistent CFP for use in future ingestive behavior research.

## 1 Introduction

Conditioned flavor preference (CFP) learning is a form of associative learning commonly used in ingestive behavior research (Myers and Sclafani, 2006, Sclafani et al., 1993, Sclafani and Ackroff, 1994). In CFP learning an animal learns to associate a novel flavor (conditioned stimulus [CS+] e.g. grape Kool-Aid) with a positive orosensory or postingestive effect (unconditioned stimulus [US] e.g. sweet taste, or calories from glucose). When tested later in a 2-bottle preference test for the CS+ versus a previously unconditioned flavor (CS- e.g. cherry Kool-Aid), without the US present, the animal shows a preference for the CS+.

CFP is significant as a model of food learning, rivaling conditioned taste aversion learning in its rapid acquisition and persistence. It is also of ecological and public health relevance, as human dietary preferences based on flavor are strong, established early in life, and may contribute to obesity and oral addictions (Beauchamp and Mennella, 2009, Beauchamp and Mennella, 2011). Because of its robustness, and the ease of acquisition, CFP may serve as a model for the neurological analysis of preference comparable to conditioned place preference or pair bonding.

The two most common methods that have been used for the study of CFP learning are pairing oral intake of the CS+ with intragastric infusion of nutrients and oral intake of nutrients or sweeteners. Oral conditioning takes advantage of both taste reinforcement and postingestive reinforcement (if calories are present). To separate out the contribution of taste reinforcement and postingestive reinforcement, some studies induce CFP using saccharin as a non-caloric tastant, or employ fructose as the US, which has been found to not strongly mediate postingestive conditioning (Ackroff et al., 2001, Golden and Houpt, 2007, Risco and Mediavilla, 2014, Touzani and Sclafani, 2005, Baker et al., 2003, Baker et al., 2004, Bernal et al., 2009, Bernal et al., 2008). Sclafani and others have used an elegant “electronic esophagus” preparation (Elizalde and Sclafani, 1990), in which oral intake of a non-nutritive solution is paired in realtime via lickometer with intragastric infusion of a nutrient US, e.g. glucose (Sclafani et al., 1993, Sclafani, 2004, Touzani et al., 2010, Myers, 2007, Perez et al., 1998, Sclafani and Ackroff, 2006, Wald and Myers, 2015). This allows orosensory and postingestive stimuli to be manipulated independently, and has been crucial to isolating the determinants of the glucose US (Zukerman et al., 2013, Sclafani and Ackroff, 2015, Sclafani et al., 2016).

In the present study we sought to establish a model of robust long-lasting CFP learning with a relatively quick and simple conditioning procedure that could be used in future research (Kimbrough et al., 2011). To do so we took advantage of flavors mixed with glucose to be consumed orally and used a conditioning procedure of 48-h to create a long-lasting preference. As conditioning controls, we established that rats did not show a preference between the two Kool-Aid flavors, and confirmed the requirement for a caloric US. We tested the duration and strength of the preference over the course of 14 days of extinction, and then again for long-term retention 48 days after conditioning.

## 2 Materials and Methods

### 2.1 Animals

Adult male Sprague–Dawley rats (300–450g, Charles River Laboratories, Wilmington, MA) were individually housed under a 12-h light–12-h dark cycle (lights on 07:00) at 25 °C. Throughout the experiments, rats had free access to Purina rodent chow and distilled water or Kool-Aid solutions as noted below. All experiments were approved by the Florida State University institutional animal care and use committee.

### 2.2 Experiment 1: Unconditioned intake of grape vs. cherry Kool-Aid

We tested unconditioned preferences of grape vs. cherry Kool-Aid (Kraft Foods, Northfield, IL) in rats in order to determine if rats show a preference for one flavor over the other. Two solutions were used in this experiment 0.05% unsweetened grape Kool-Aid mixed with 0.05% saccharin and 0.05% unsweetened cherry Kool-Aid mixed with 0.05% saccharin. Naïve rats (n=8) were given 24-h, 2-bottle preference tests with both grape and cherry Kool-Aid solutions. Intake was recorded by bottle weight and solutions replaced every 24 h of 2-bottle testing, and the position of each bottle was swapped daily to observe side biases. The testing period lasted for 14 days.

### 2.3 Experiment 2: Effect of CS Pre-Exposure

We tested the effect of pre-exposing rats to a flavor (CS+) prior to preference testing with the same flavor (CS+) and another non-pre-exposed flavor (CS-). Two solutions were used in this experiment: 0.05% grape Kool-Aid mixed with 0.05% saccharin and 0.05% cherry Kool-Aid mixed with 0.05% saccharin. Naïve rats (n=8) were given 48-h 2-bottle access to either grape or cherry Kool-Aid solution (CS+; counterbalanced for each flavor) and water. The bottle positions of the CS+ and water were swapped after 24 h.

Following pre-exposure 24-h 2-bottle testing was performed with both the pre-exposed CS+ flavor and the novel CS- flavor. Intake was recorded every 24 h of 2-bottle testing and the position of each bottle was swapped daily. The testing period lasted for 14 days.

### 2.4 Experiment 3: 48-h glucose conditioning

After testing for a preference being established by pre-exposure to a novel flavor for 48-h (experiment 2), we tested whether 48-h conditioning with a CS+ flavor mixed with a palatable tastant (glucose) would result in a long lasting preference. During conditioning naïve rats (n=8) were given 48-h 2-bottle access to a CS+ flavor (0.05% grape or cherry Kool-Aid [counterbalanced for flavor] mixed with 8% glucose and 0.05% saccharin) and water. The position of the CS+ and water bottles were swapped after 24 h.

Following conditioning rats were given 24-h, 2-bottle preference tests of the CS+ flavor and the unconditioned flavor (grape or cherry; CS-) with no glucose in either solution (both solutions still contained 0.05% Kool-Aid and 0.05% saccharin). Intake was measured every 24 h and the position of each bottle was swapped daily. The testing period lasted for 14 days. Up until this point experiments 2 and 3 were identical except for the presence of glucose during conditioning in experiment 3.

After the 14 days of extinction testing, rats were maintained on rodent chow and water without Kool-Aid access. At 48 days after conditioning (34 days after the last extinction test), rats were tested again with CS+/saccharin vs. CS-/saccharin, following the same 2-bottle procedure as above for 2 consecutive days with the bottle position swapped on the second day.

### 2.5 Statistics

We calculated Kool-Aid preferences as percent of total intake: Kool-Aid / (Kool-Aid and water), or CS+ / (CS+ and CS-). We ran two-way repeated measures ANOVAs on each of the 14-day 2-bottle preference tests in each experiment. We ran post-hoc Student Newman Kuels (SNK) on all 14-day preference tests. In experiment 3, a t-test of the 2-day average 48 days after conditioning was done to test for significance.

## 3 Results

### 3.1 Experiment 1: Unconditioned intake of grape vs. cherry Kool-Aid

We tested if rats had an unconditioned preference for either grape or cherry Kool-Aid over the other flavor. The average daily intake per rat across the 14 days of 2-bottle testing was 35.3±6.3g for grape Kool-Aid/saccharin and 22.6±3.4g for cherry Kool-Aid/saccharin.

Two-way repeated measures ANOVA revealed no effect of groups (cherry vs. grape), but a significant effect of days (F(2,13)=5.081, p<0.05) and a significant interaction between days and group (F(12,13)=5.446, p<0.05) (see Figure 1A). Post-hoc SNK analysis revealed significant differences (p<0.05) between cherry and grape intake on days 2,6,8,10, and 14.

**Figure 1:**
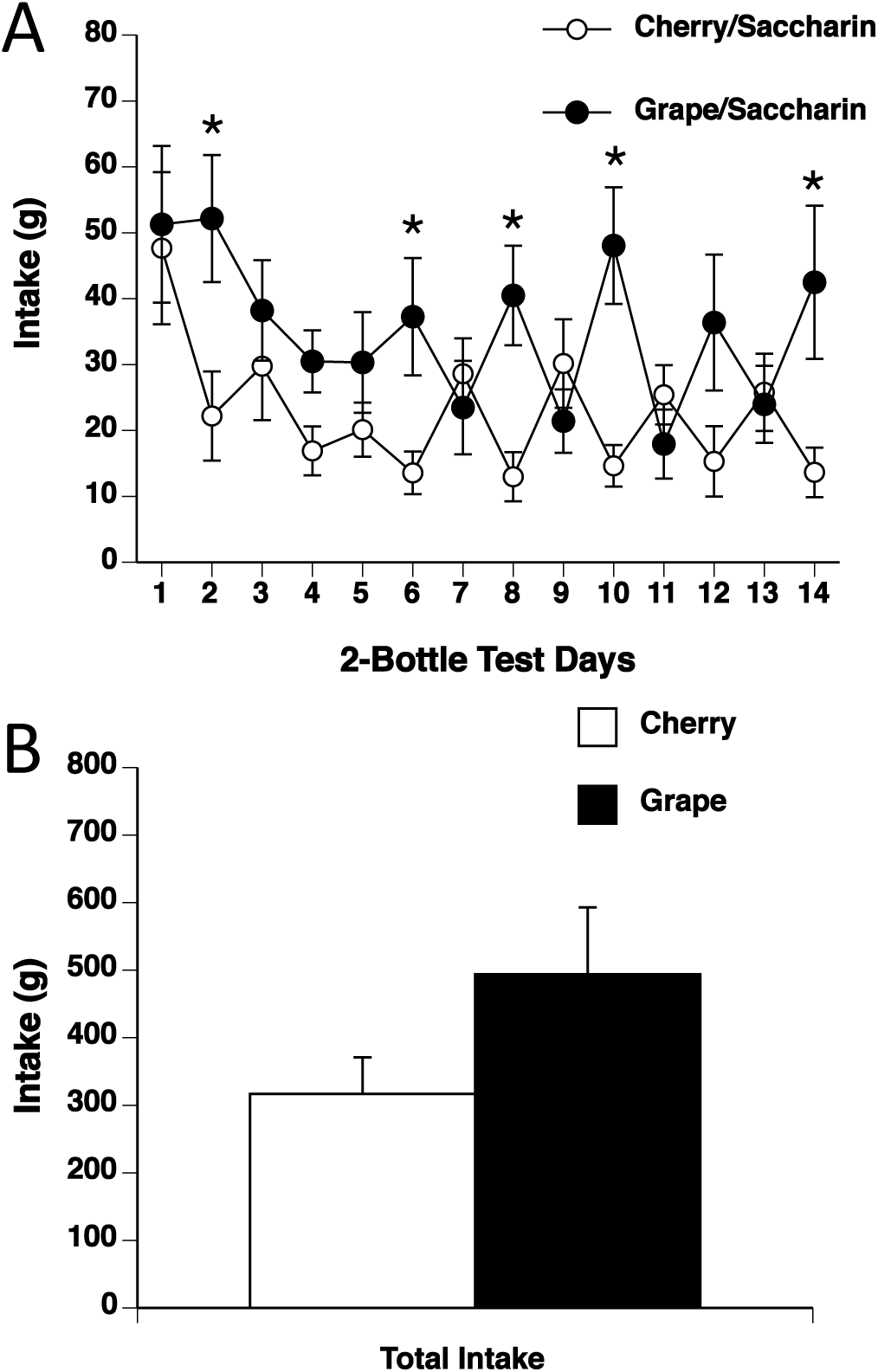
Unconditioned grape and cherry Kool-Aid preferences. (A) Intake over 14 days of two-bottle preference testing of Kool-Aid flavors grape/saccharin (black circles) and cherry/saccharin (white circles). Rats consumed significantly more grape Kool-Aid on days 2, 6, 8, 10, and 14. (B) Total intake of Kool-Aid Flavors grape/saccharin (black bars) and cherry/saccharin (white bars) across the 14 days of two-bottle preference testing. Rats did not drink significantly more of either flavor of Kool-Aid over the 14 days. * p<.05 grape *vs*. cherry.

There was no significant difference between total intake of grape and cherry Kool-Aid across the 14 days (see Figure 1B). Thus, rats showed a slight preference for grape over cherry Kool-Aid on some days, but the total intakes are not different.

### 3.2 Experiment 2: Effect of CS Pre-Exposure

We also tested whether rats would form a preference for one flavor (grape or cherry Kool-Aid) over the other after a 48-h pre-exposure to one flavor (CS+). There was no difference in intake between rats given grape or cherry Kool-Aid during the 48-h pre-exposure (see Figure 2A). The average daily intake per rat across the 14 days of 2-bottle testing was 27.2±2.6g for CS+/saccharin and 26.6±3.6g for CS-/saccharin. Two-way repeated measures ANOVA revealed no effect of pre-exposure group (cherry vs. grape), no effect of days, and no interaction. However, by t-test CS-/saccharin intake was greater than CS+/saccharin intake on day 1 (see Figure 2B). There was no difference between total 14-day intake of CS+ and CS- flavors (see Figure 2C). Thus, CS+ pre-exposure does not seem to cause a lasting preference for either the CS+ or the CS-.

**Figure 2:**
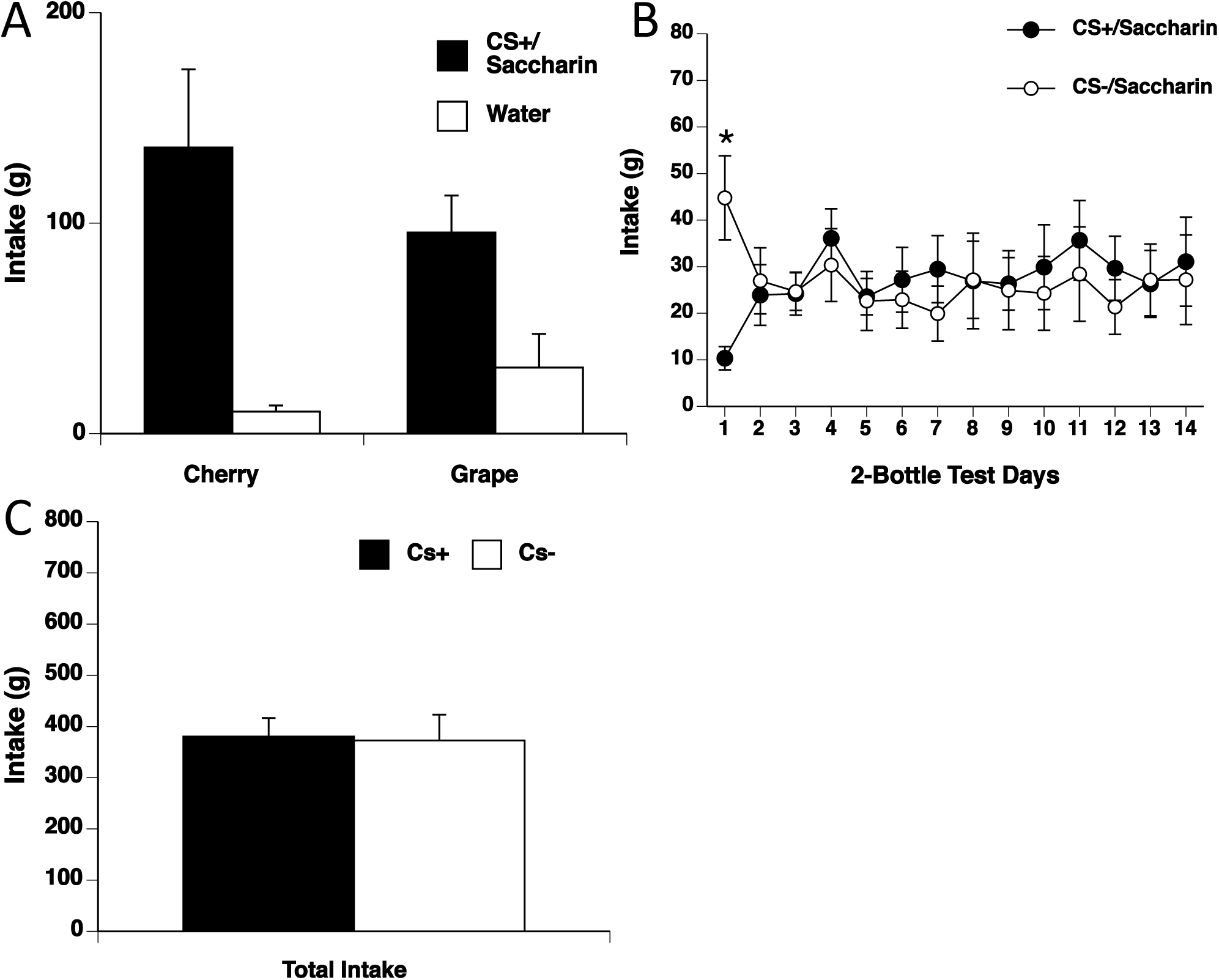
48-h pre-exposure to a CS flavor without glucose. (A) 48-h total intake during pre-exposure to CS+/saccharin. Rats drank more CS+ (black bars; grape or cherry) than water (white bars) during the 48-h pre-exposure. (B) Intake over 14 days of two-bottle preference testing of CS+/saccharin (black circles) and CS-/saccharin (white circles). Rats did not drink more of either the CS+ or CS- on any day except day one, which rats drank significantly more CS- than CS+. (C) Total intake of CS+/saccharin (black bars) and CS-/saccharin (white bars) across the 14 days of two-bottle preference testing. Rats did not differ in amount of CS+ and CS- consumed over the 14 days after 48-h pre-exposure to the CS+ flavor. Thus, pre-exposure to a CS+ flavor alone did not lead to a CFP. * p<.05 CS+ *vs*. CS-.

### 3.3 Experiment 3: 48 h glucose conditioning

We conditioned rats with a CS+ Kool-Aid flavor mixed with 8% glucose for 48-h, and then determined the strength and persistence of the acquired CFP. Glucose conditioned rats consumed 301.9±17.3g CS+/glucose and 7.5±0.6g water during conditioning. There was no significant difference between grape or cherry intake during conditioning (see Figure 3A).

**Figure 3:**
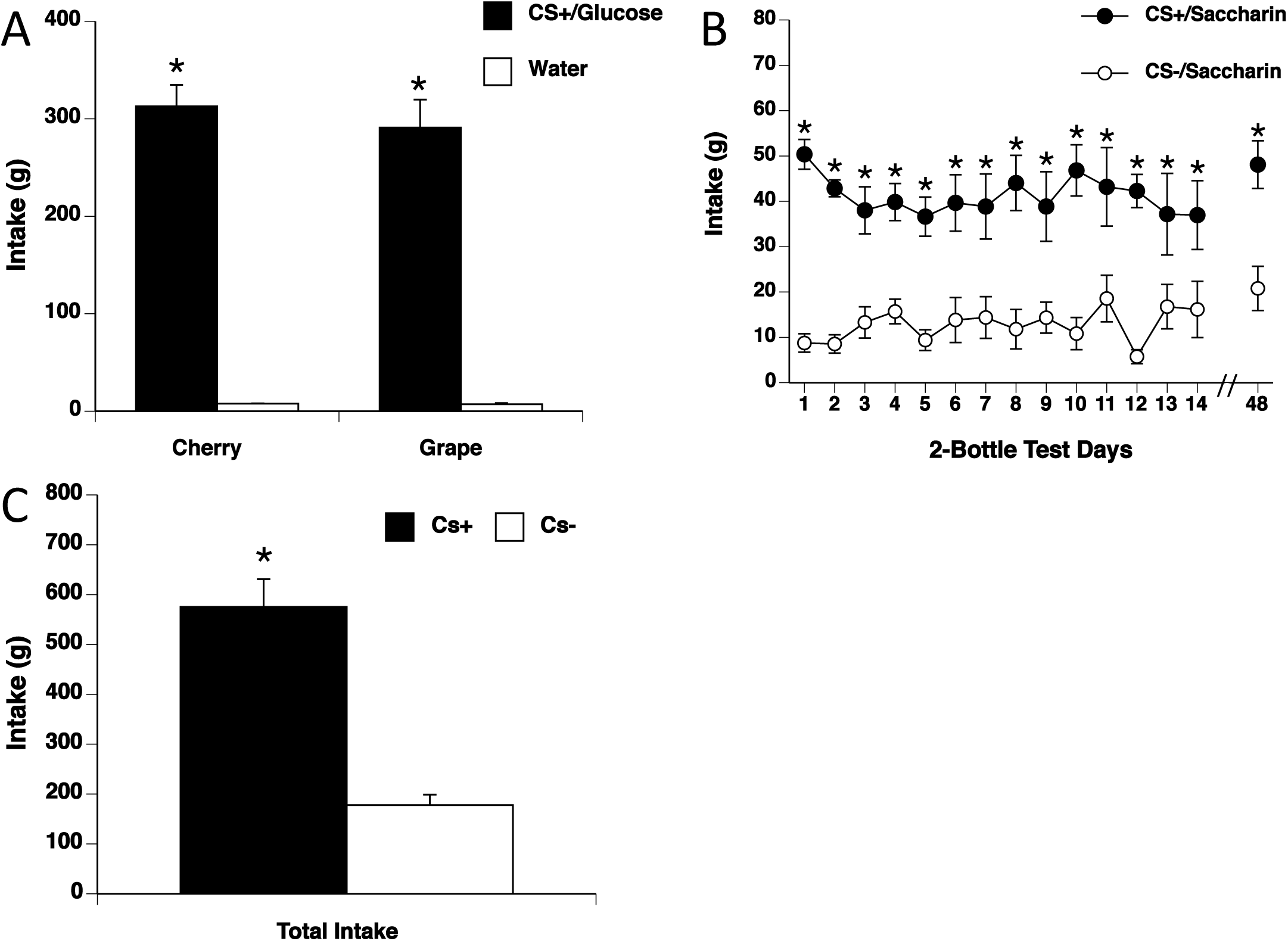
48-h conditioning with CS+ flavor and glucose. (A) 48-h total intake during conditioning with CS+/8% glucose (grape and cherry). Rats receiving both grape and cherry as the CS+ consumed significantly more CS+/glucose (black bars) during the 48-h conditioning than water (white bars). (B) CS+ and CS- intake during 14 days of two-bottle preferences tests and average 48-h intake on day 48 post conditioning. Rats showed a significantly greater intake of CS+/saccharin (black circles) than CS-/saccharin (white circles) at each day, including 48 days later. (C) Total intake of CS+/saccharin (black bars) and CS-/saccharin (white bars) across the first 14 days of two-bottle preference tests. Rats consumed significantly more CS+/saccharin than CS- saccharin throughout the 14 days of two-bottle preference testing. Thus, rats formed a robust, long lasting and resistant to extinction CFP after 48-h of conditioning with CS+/glucose. * p<.05 CS+ *vs*. CS- or CS+ *vs.* water.

After conditioning rats had an initial intake 24-h intake of 50.4±3.3g CS+/saccharin and 8.8±2.0g CS-/saccharin, and a CS+ preference of 86±3%. The preference persisted across 14-days of 2-bottle extinction (preference of 69±11% and intakes of 37.0±7.6g CS+/saccharin vs. 16.2±6.2g CS-/saccharin on day 14 of extinction). Two-way repeated measures ANOVA revealed CS+ intake to be significantly greater than CS- intake (F(1,13)=45.30, p<0.05, CS+ vs. CS-), no effect of days, and no interaction. Post-hoc SNK analysis revealed significant differences (p<0.05) between CS+ and CS- intake across all 14 days (see Figure 3B). There was a significant difference (p<0.05) between total 14-day intake of CS+ and CS- (see Figure 3C).

When rats were tested 48 days after conditioning (34 days after the last 2-bottle extinction test), they showed an average 48-h CS+ preference of 70±7% and average 48-h intake of 48.1±5.3g CS+/saccharin vs. 20.8±4.9g CS-/saccharin for the 2 test days. T-test of CS+ vs. CS- for the 48-h average intake showed that intake of the CS+ flavor was still significantly higher than CS- intake (see Figure 3C, p <.05).

Thus, rats form a strong preference for a CS+ Kool-Aid flavor after 48-h conditioning with glucose. The preference formed is long lasting and persists over a month after conditioning

## 4 Discussion

The primary goal of this study was to establish a method of CFP learning that would produce a strong and long lasting preference in rats. In experiment 1 we tested if rats showed an unconditioned preference for either grape or cherry Kool-Aid in our hands. As expected we found rats did not show a larger or significant preference for either flavor of Kool-Aid when tested without prior conditioning.

In experiment 2 we tested if pre-exposure to one flavor (CS+) mimicking the 48-h conditioning period but without the reinforcement of glucose would lead to a preference. We found that pre-exposure to the CS+ did not lead to a significant preference for the CS+ and actually produced a preference for the CS- on the first day of 2-bottle testing that did not persist to additional days. The initial preference for the unconditioned flavor was not expected, however it may be due to sensory specific satiety of the CS+ given there was no reinforcement during pre-exposure. Sensory specific satiety is the decreased intake of a previously ingested food when tested versus a novel food and has been seen in both humans (Havermans et al., 2009, Rolls et al., 1981) and rats (Berridge, 1991).

In experiment 3 we found that 48-h conditioning with Kool-Aid reinforced by oral glucose produced a strong, long-lasting CFP, which was resistant to extinction. The CFP produced persisted not only though 14 days of 24-h extinction testing, but additionally for 48 days post-conditioning (including extinction). We can speculate that if testing had continued beyond this point that the CFP would have lasted even longer.

In experiment 2 rats did not show a preference for a pre-exposed, or familiar flavor, indicating familiarity alone is not enough to produce a preference. However, we cannot conclude with complete certainty that familiarity did not play a role in enhancing or maintaining the learned CFP in experiment 3. Thus, there is the possibility that the taste-nutrient reinforcement during conditioning did not establish a preference without other factors. Instead a combinatorial effect of taste-nutrient reinforcement during conditioning and familiarity with the flavor may have worked together to produce a robust preference for the CS+.

Another CFP paradigm that does not make use of the intragastic infusion preparation has been reported previously (Warwick and Weingarten, 1994). In the paradigm described by Warwick and Weingarten (1994) the authors present an elegant method of establishing a CFP that is able to dissociate nutrient and taste components CFP learning, something the paradigm in the present report is unable to do. Instead the current paradigm offers a method to establish a robust, long lasting CFP in a relatively short conditioning time window (48-h).

Our conditioning parameters and observed results are in line with previous studies. A CFP can be established in as little as a single 10–30 min session when a flavor is paired with gastric infusions of nutrients (Ackroff et al., 2009, Myers, 2007). A few studies have looked at extended extinction testing and shown preferences that have lasted from at least 13 days up to 40 days, and preferences can persist 30 days after extinction testing ends (Myers, 2007, Kimbrough et al., 2011, Elizalde and Sclafani, 1990, Drucker et al., 1994).

In conclusion, we have demonstrated a simple and rapid procedure to induce a robust and persistent flavor preference in a discrete period of time. These data confirm and extend observations made about the persistence and robustness of CFP learning.

## Acknowledgements

Supported by NIDCD T32-000044 (AK).

